# Mitochondrial uncouplers impair human sperm motility without altering ATP content

**DOI:** 10.1101/2022.08.04.502752

**Authors:** Will M. Skinner, Natalie T. Petersen, Bret Unger, Shaogeng Tang, Emiliano Tabarsi, Julianna Lamm, Liza Jalalian, James Smith, Ambre M. Bertholet, Ke Xu, Yuriy Kirichok, Polina V. Lishko

## Abstract

Sperm motility is necessary for successful fertilization, but there remains controversy about whether human sperm motility is primarily powered by glycolysis or oxidative phosphorylation. To evaluate the plausibility of reducing human sperm mitochondrial ATP production as an avenue for contraceptive development, we treated human sperm with small-molecule mitochondrial uncouplers, which reduce mitochondrial membrane potential by inducing passive proton flow, and evaluated the effects on a variety of physiological processes that are critical for fertilization. We also sought to clarify the subcellular localization of Adenosine Nucleotide Translocator 4 (ANT4), a gamete-specific protein that has been suggested as a contraceptive target. We determined that ANT4 is mitochondrially localized, that induced mitochondrial uncoupling can be partially mediated by the ANT family, and that two uncouplers, Niclosamide Ethanolamine and BAM15, significantly decreased sperm progressive motility. However, these uncouplers did not reduce sperm ATP content or impair other physiological processes, implying that human sperm can rely on glycolysis for ATP production in the absence of functional mitochondria. Thus, since certain mitochondrial uncouplers impair motility through ATP-independent mechanisms, they could be useful ingredients in on-demand, vaginally-applied contraceptives. However, systemically delivered contraceptives that target sperm mitochondria to reduce their ATP production would need to be paired with sperm-specific glycolysis inhibitors.

**Significance Statement:** Development of novel contraceptives is critical, since half of all pregnancies are still unplanned, even in developed countries. This high unplanned pregnancy rate contributes to a wide variety of social, environmental, and ecological problems. Impairing human sperm is a way to develop male and unisex contraceptives, but much remains unknown about these unique cells. Here we settle a long-running debate about human sperm metabolism, finding that human sperm can maintain their ATP levels without mitochondrial oxidative phosphorylation. This finding will help focus future contraceptive development efforts. We also identify the potential use of an FDA-approved compound (Niclosamide) as a motility-impairing ingredient in spermicides and correct the misunderstood subcellular localization of an existing contraceptive target, Adenosine Nucleotide Translocator 4.

## Introduction

Human sperm make one of the most remarkable and challenging journeys of any cell in the human body. To reach the egg, the mature sperm must traverse a distance thousands of times greater than their cell length, while surviving environments that differ greatly in viscosity, pH, and oxygen and sugar concentrations (1,2). This journey requires vigorous flagellar motility (3), and later asymmetrical “hyperactivated” flagellar bending (4), both of which require large amounts of ATP. However, each human sperm possesses less than a fifty mitochondria (5), less than almost any other metabolically active cell. Therefore, there has been persistent debate about the role of mitochondria and oxidative phosphorylation in sperm physiology, with metabolic studies showing that glycolysis is sufficient for human sperm ATP production (6), but physiological and clinical studies showing that proper mitochondrial functionality is correlated with high fertility in humans (7–10). Additionally, sperm contain many unique proteins and protein isoforms, including metabolic enzymes (11), membrane transporters (12,13) and receptors (14), and structural proteins (15). Many of these unique proteins have been appealing targets for development of contraceptives that could disrupt sperm bioenergetics.

One of these unique transporter isoforms is Adenosine Nucleotide Translocator 4 (ANT4), which is gamete-specific (16) and required for spermatogenesis (17), but has an unclear role in mature human sperm cells. Since its discovery, ANT4 has been seen as an attractive candidate for the development of a male contraceptive (17–20). Efforts have been made to develop specific ANT4 inhibitors, which were only able to achieve modest specificity (19,20). Somatic ANT isoforms (ANT1, 2, and 3 in humans) are essential for transporting ATP and ADP across the inner mitochondrial membrane (IMM) and have recently been found to be responsible for mediating mild mitochondrial proton leak (or mitochondrial “uncoupling”) across the IMM in somatic, non-thermogenic tissues (21). It is also possible to activate ANT-mediated proton leak using exogenous small molecule uncouplers (22). Therefore, we hypothesized that instead of developing specific ANT4 inhibitors, it may be possible to develop novel small molecule uncoupler contraceptives that act by specifically activating proton leak through ANT4, thereby decreasing sperm mitochondrial membrane potential (MMP), ATP production, and ultimately functionality. To pursue this hypothesis, we first needed to validate the idea that breakdown of sperm MMP leads to functional dysregulations that could impair fertility. To do so, we began by testing the physiological effects of various small molecule uncouplers on human sperm cells.

## Results

### Representative small molecule uncouplers decrease human sperm mitochondrial membrane potential and impair basal motility

Four representative small molecule uncouplers were chosen to assess the effects of this class of compounds on human sperm physiology: 2, 4, Dinitrophenol (DNP); carbonyl cyanide p-trifluoromethoxyphenylhydrazone (FCCP); BAM 15 (N5,N6-bis(2-Fluorophenyl)-[1,2,5]oxadiazolo[3,4-b]pyrazine-5,6-diamine); and Niclosamide Ethanolamine (NEN) (Fig. 1A). Live human sperm cells were treated with varying concentrations of each compound for 30 minutes at 37°C in noncapacitating media at pH 7.4, after which cells were stained with MitoTracker Red CMX-Ros and mitochondrial membrane potential (MMP) was qualitatively assessed using flow cytometry (Fig. 1B) and fluorescence microscopy (Fig. 1D). FCCP, BAM15, and NEN were all able to dose-dependently reduce midpiece fluorescence and therefore MMP, while DNP was unable to reduce midpiece fluorescence at concentrations up to 300 μM at this pH level (Fig. S1A). The DMSO and Ethanol vehicles with which compounds were delivered did not have a significant effect on mitochondrial fluorescence (Fig. S1A) and none of the compounds decreased sperm viability (Fig. S1D). NEN and BAM15 reduced midpiece fluorescence values more strongly than FCCP, had lower IC_50_ values, and are known to have more favorable toxicity profiles (23,24), and so further experiments focused on these two compounds.

**Figure 1:**
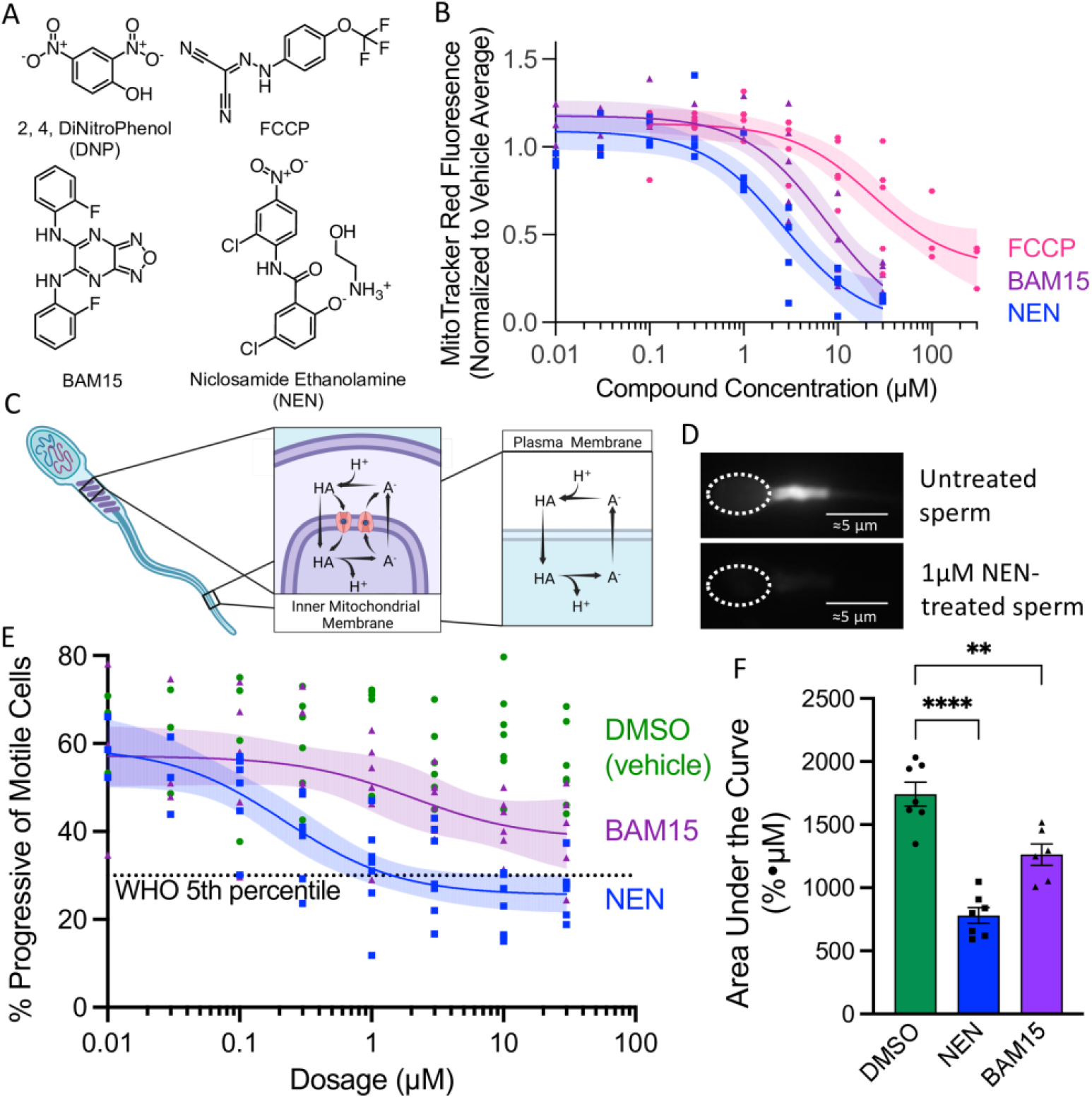
Representative small-molecule uncouplers dose-dependently reduce sperm mitochondrial membrane potential and basal motility. **A)** Chemical structures of representative small molecule uncouplers used. **B)** Effects of SMUs on human sperm mitochondrial potential as measured by MitoTracker Red fluorescence, normalized to average fluorescence of relevant vehicle. Nonlinear least squares regression performed using an unweighted [inhibitor] vs. response three parameter model (Hill Slope = -1.0), identifying “unstable” fits. Shaded areas indicate 95% confidence bands for true location of each nonlinear fit curve. The nonlinear regression fit had an Adjusted R^2^ value of .8409 for NEN, .8179 for BAM15, and .7143 for FCCP. Adjusted R^2^ value for DNP was less than .1 and so curve is not shown or reported. Asymmetrical 95% CI intervals of IC_50_ for NEN: 1.415 μM to 5.380 μM; BAM15: 3.827 μM to 17.03 μM; FCCP: 9.148 μM to 64.05 μM. **C)** Model showing the possible mechanisms of action of small molecule uncouplers in the sperm mitochondria and plasma membrane. **D)** Representative fluorescence micrographs of sperm cells treated with MitoTracker Red CMX-Ros in the absence (top) and presence (bottom) of 1 μM NEN. **E)** Effects of SMU on human sperm progressive motility as measured by Computer-Aided Sperm Analysis. Data points represent the average of ≥500 sperm cells. Nonlinear least squares regression performed and displayed as in B. The nonlinear regression fit had an Adjusted R^2^ value of .6323 for NEN, .2722 for BAM15, and <.1 for DMSO. Asymmetrical 95% CI interval for NEN: 0.08256 μM to 0.6284 μM. Asymmetrical 95% CI interval for BAM15: 0.1264 μM to 78.30 μM. **F)** Area Under the Curve quantification for graph E. Error bars represent SEM, and each data point represents the area under the curve for all concentrations of one biological replicate. Statistical significance calculated using Dunnett’s multiple comparisons test.

Next, we assessed the effects of NEN and BAM15 on human sperm motility, using computer-aided sperm analysis (CASA). Both NEN and BAM15 reduced the percent of sperm showing progressive motility (Fig. 1E,F) and NEN reduced the average curvilinear velocity (Fig. S2A); NEN reduced sperm progressive motility percent below the 5^th^ percentile of fertile men (25) and showed a sub-micromolar IC_50_.

### Adenosine Nucleotide Translocators (ANTs) partially mediate NEN-induced mitochondrial uncoupling, and ANT4 is localized to the human sperm mitochondria

In somatic, non-thermogenic tissues of the body, Adenosine Nucleotide Translocator (ANT) proteins are known to mediate mild mitochondrial uncoupling under physiological conditions and in the presence of exogenous uncouplers by inducing H^+^ influx across the inner mitochondrial membrane (21,22). Since sperm cells possess a unique isoform of the family known as ANT4 (26), we investigated whether NEN-induced uncoupling is also mediated by the ANT family of proteins. To do this, we performed mitochondrial patch-clamp electrophysiology, through which it is possible to directly measure current across the inner mitochondrial membrane. Indeed, in mouse heart mitochondria, which abundantly express ANT1 (27), 500 nM NEN induced a strong inward H^+^ current at physiological pH and negative voltage. This H^+^ current was significantly reduced both in the presence of 1 μM Carboxyatractyloside (CATR), a known ANT inhibitor (Fig. 2B, C), and in heart mitochondria from ANT1-knockout mice (Fig. 2B, D), indicating that the NEN-induced current is partially mediated by ANT proteins.

**Figure 2:**
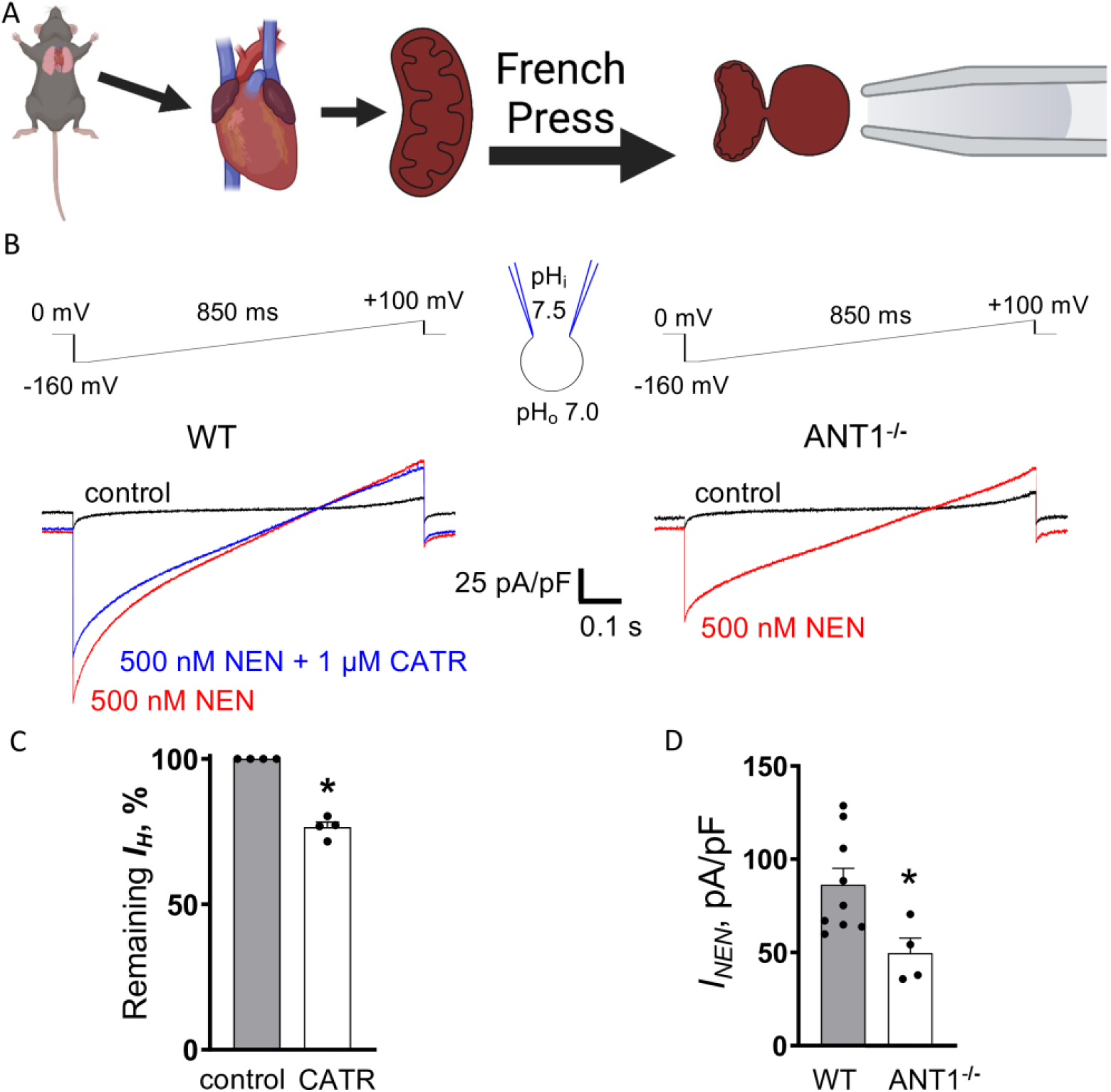
NEN induces partially ANT-mediated H^+^ current in mouse heart mitoplasts. **A)** Diagram of mouse heart mitoplast isolation and preparation for whole-organelle inner membrane electrophysiology. **B)** Representative whole-organelle electrophysiological traces of H^+^ current density in response to voltage ramp protocols performed on mouse heart mitoplasts. Left, Wildtype; Right, ANT1 knockout. **C)** Quantification of NEN-induced H^+^ current across mitochondrial inner membrane before and after application of 1 μM CATR, a known ANT inhibitor. Significances calculated by Student’s unpaired t-test. **D)** Quantification of H^+^ current density induced by 500 nM NEN in wildtype and ANT1-knockout mouse heart mitoplasts.

Given this possibility of ANT-mediated uncoupling occurring in sperm, and the presence of a unique isoform (ANT4) in sperm cells, we investigated whether ANT4 is indeed expressed in the sperm mitochondria. Other investigators have proposed that ANT4 is expressed throughout the sperm flagellum, and hypothesized that it may function in collaboration with glycolytic enzymes to traffic ATP throughout the sperm flagellum (28). However, all other ANT isoforms are expressed exclusively on the inner mitochondrial membrane, so a flagellar localization would be surprising. Using immunocytochemistry and confocal microscopy, we found that when excess primary antibody is used, some ANT4 staining is indeed seen throughout the flagellum (Fig. S3F), but when primary antibody concentration is titrated down to an optimal concentration, that staining is restricted to the midpiece in a similar distribution to that of the inner mitochondrial membrane protein Cytochrome c oxidase (COXIV) (Fig. 3A-C). This ANT4 staining is not seen in non-permeabilized sperm (Fig. S3D), suggesting an exclusively mitochondrial localization consistent with other ANT isoforms. To confirm this finding, we conducted stochastic optical reconstruction microscopy (STORM) on human sperm cells similarly costained for COXIV and ANT4 (Fig. 3D-F, top) and found that upon virtual cross-sectioning of the midpiece, their distributions formed rings of similar diameter and thickness (Fig. 3D-F, bottom), consistent with mitochondrial localization.

**Figure 3:**
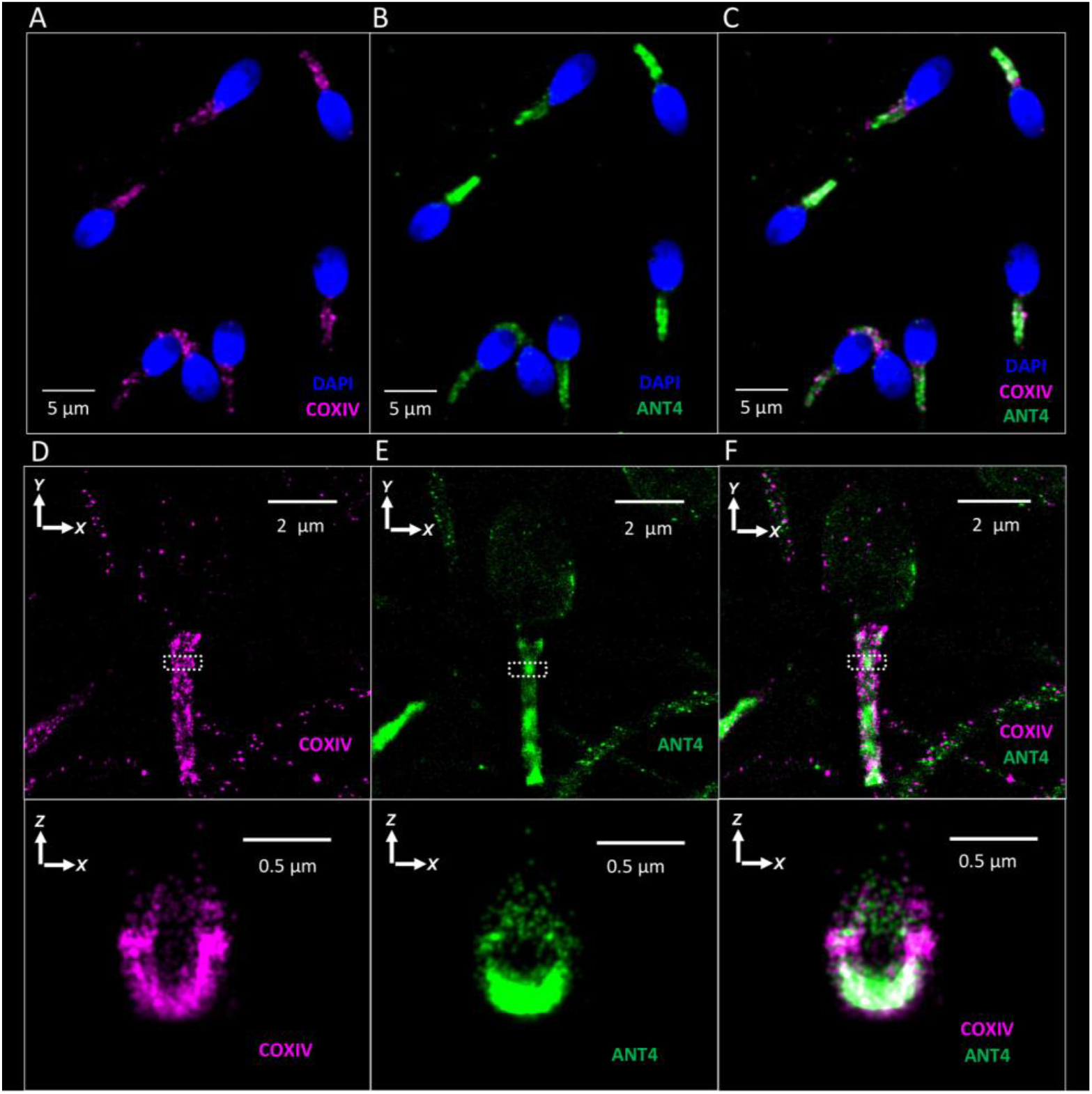
ANT4 is localized to the human sperm mitochondria. **A-C)** Confocal microscopy of human sperm immunocytochemistry showing the localization of COXIV (magenta), ANT4 (green), and DAPI (blue) **D-F)** Stochastic optical reconstruction microscopy (STORM) of human sperm cells showing the localization of COXIV (magenta) and ANT4 (green). Top panels: overhead view (XY plane) of sperm cells. Bottom panels: virtual cross section (XZ plane) of midpiece area indicated in top panels by white dotted box.

### Small molecule uncouplers do not reduce human sperm ATP content, and NEN induces significant proton current across the sperm plasma membrane

To investigate the mechanism of action of NEN and BAM15’s effect on sperm basal motility (Fig. 1E, F) we assessed their effects on sperm ATP levels and found that neither compound had any effect on ATP concentrations in noncapacitated (Fig. 4A) or capacitated (Fig. 4B) spermatozoa. Consistent with this finding, we observed that NEN and BAM15 had no effect on other ATP-dependent human sperm physiological processes, including the acrosome reaction, a necessary pre-fertilization exocytotic event (29,30) that exposes key fertilization receptors (31,32) (Fig. S4A – C), and human sperm hyperactivated motility (33) (Fig. S4D, E). NEN also did not impair *in vitro* fertilization between mouse sperm and eggs (Fig. S4F, G).

**Figure 4:**
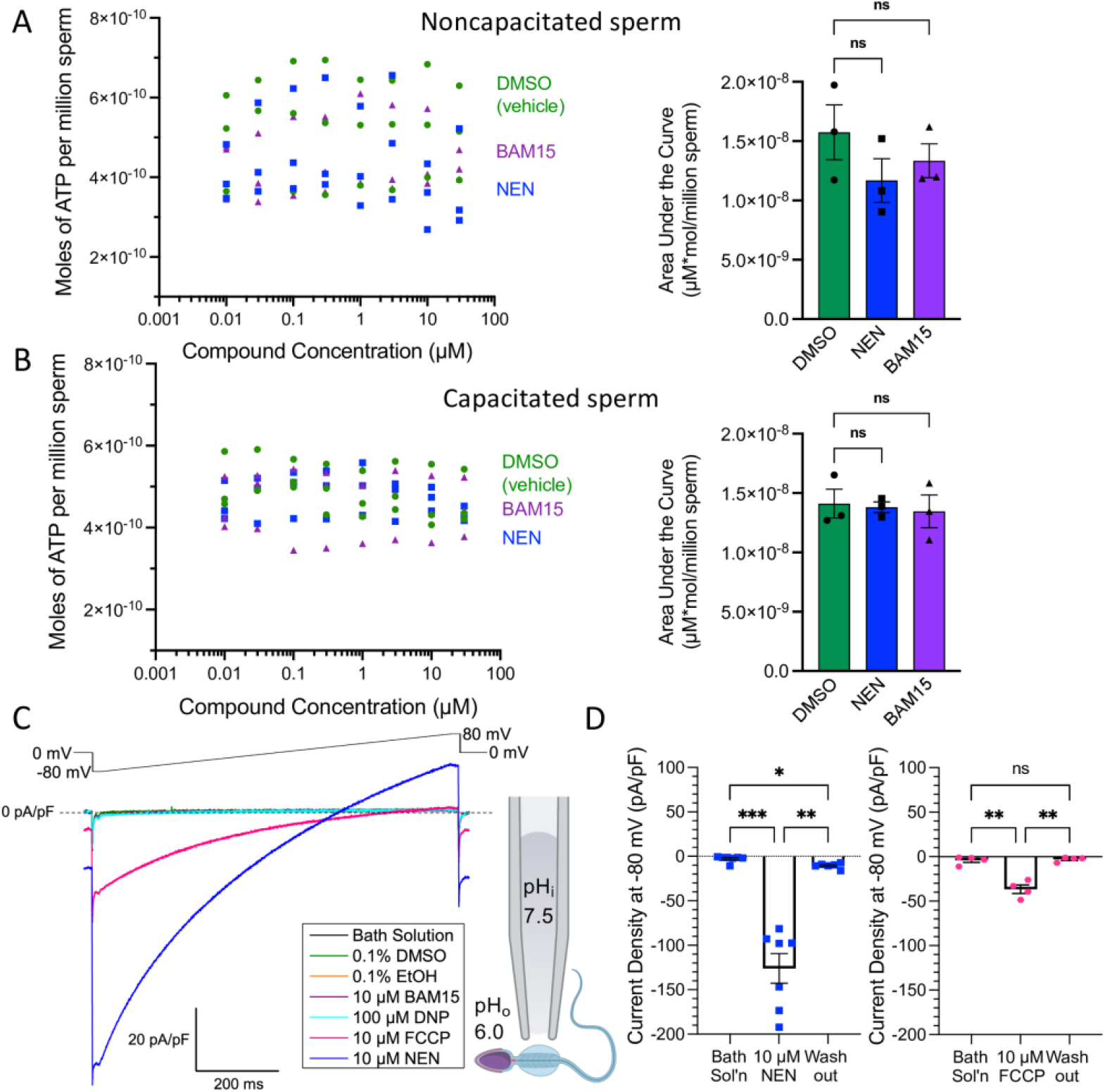
Effects of small molecule uncouplers on human sperm ATP production and sperm membrane proton current. **A)** Left, quantification of ATP content of non-capacitated human sperm treated with NEN, BAM15, and DMSO vehicle. Right, area under the curve analysis of the plot to the left. **B)** Left, quantification of ATP content of human sperm capacitated in the presence of NEN, BAM15, and DMSO vehicle. In both A and B, nonlinear regressions not shown, because all adjusted R^2^ values were <.1. Outliers in both A and B tested using a ROUT coefficient of 1%, with no outliers detected. Error bars in all plots represent SEM. Statistical significance in AUC analysis calculated using Dunnett’s multiple comparisons test. **C)** Left, Representative electrophysiological traces showing the effect of vehicles and uncouplers on human sperm membrane H^+^ current density. Right, diagram of sperm electrophysiology setup. Cytoplasmic droplet is magnified for visibility. **D)** Quantification of maximal current densities in the presence of 10μM NEN (left) and FCCP (right), at a voltage of -80mV. Error bars represent SEM, and significance calculated using Tukey’s Multiple Comparisons Test.

Since the uncouplers’ negative effects on sperm basal motility were not mediated by a reduced amount of cellular ATP, to further investigate their mechanism of action we assessed the effects of each compound on sperm plasma membrane H^+^-ion current using whole-cell patch clamp electrophysiology. In this assay, 10 μM NEN and 10 μM FCCP induced a significant inward transmembrane H^+^-ion current at negative voltages (Fig. 4C, D), while 10 μM BAM15, 100 μM DNP, and the Ethanol and DMSO vehicles induced no discernable H^+^ current (Fig. 4C, S5A). The maximum current density of 10 μM FCCP (Fig. 4D) and 100 μM DNP (Fig. S5A) was consistent with recently reported values in the plasma membrane of HEK293 cells (22). Additionally, the current density induced by 10 μM NEN across the sperm plasma membrane (Fig. 4D) was only roughly 1.5 times greater than the current density caused by a 20x lower concentration of NEN in the inner mitochondrial membrane (IMM) (Fig. 2D). This buttresses the idea that part of NEN’s current is mediated by ANT proteins located exclusively in the IMM, which increase the IMM’s permeability to NEN-induced current. Although NEN took significantly longer than FCCP to reach its maximal transmembrane current, their maximal rates of change of current density were not significantly different, indicating that their mechanisms of action may be similar (Fig. S5B, C).

## Discussion

This study sheds light on several key aspects of sperm energetics and physiology. There has been an ongoing debate in the field about whether oxidative phosphorylation or glycolysis is the primary source of ATP for human spermatozoa (34–36), with increasing evidence supporting the idea that glycolysis is sufficient to support many human sperm functions (6). Our data support this conclusion, with both NEN and BAM15 failing to reduce human sperm’s ATP content or ability to undergo the acrosome reaction or hyperactivated motility. Nonetheless, both uncouplers had an inhibitory effect on sperm basal motility, which in the case of NEN is likely to be related to its induction of current across the sperm plasma membrane (Fig. 4C, D), which would in turn affect many ion channels and intracellular signaling pathways that rely on a carefully moderated membrane potential for proper function. Since BAM15 is known as a more mitochondria-specific uncoupler (22,37–39) and did not induce plasma membrane current in our sperm electrophysiology assays (Fig. 4C, S5A) the mechanism of action of its more modest inhibition of basal progressive motility remains to be clarified. The fact that DNP, a known mitochondrial uncoupler (22), failed to reduce mitochondrial membrane potential may be related to DNP’s low pK_a_ (23,40) limiting its permeability into living sperm cells and/or the amount of current produced at pH 7.4, or due to the different complement of ANT isoforms present in sperm mitochondria versus skeletal muscle mitochondria (16,17,41,42), which may bind DNP with different affinities.

Although our study found that ANT4 was indeed mitochondrially localized and that the ANT family of proteins can mediate NEN-induced uncoupling, our data suggest that contraceptives that solely interfere with mature sperm oxidative phosphorylation will be unlikely to successfully inhibit key sperm functions. However, pairing an oxidative phosphorylation inhibitor with an inhibitor of a sperm-specific glycolytic enzyme, like lactate dehydrogenase C (43), might produce an effective contraceptive product. Additionally, since niclosamide has been FDA approved for use in humans with a favorable safety profile (24), and possesses low bioavailability (44), a pK_a_ close to 6 (45) (close to the vaginal pH), and significant deleterious effects on sperm motility, it could be an useful ingredient in on-demand vaginal contraceptive products as an alternative to traditional nonoxynol 9-based spermicides.

## Materials and Methods

All experiments were performed in accordance with NIH guidelines for animal research and approved by UC Berkeley Animal Care and Use Committee (AUP 2015-07-7742), with every effort made to minimize animal suffering. All described methods are consistent with the recommendations of the Panel on Euthanasia of the American Veterinary Medical Association and IACUC committee. All experimental procedures utilizing human derived samples were approved by the Committee on Human Research at the University of California, Berkeley, IRB protocol number 2013-06-5395.

### Reagents

- Niclosamide Ethanolamine was purchased from AdipoGen Life Sciences, #AG-CR1-3644
- BAM15 was purchased from Sigma, product number SML1760-5MG
- 2,4, dinitrophenol was purchased from TCI, product D0109-25G
- FCCP was purchased from abcam, product code ab120081.
- Mitotracker Red CMXRos was purchased from ThermoFisher, #M7512
- Polyclonal Antibody to human ANT4 – Novus Biologicals NBP1-89074 raised in Rabbits against human ANT4 peptide
- Mouse Monoclonal Antibody against human COX4l1 – Proteintech #66110-1-Ig
- Polyvinyl Alcohol was purchased from Fluka, #81366-100G
- A23187 was purchased from MilliporeSigma, #C7522-5MG.
- 10% Goat Serum from Invitrogen, #50-062Z
- Bovine Serum Albumin from Sigma, #A3059-500G

### Sperm Preparation

Human sperm cells were collected by masturbation from healthy donors and visually inspected for normal morphology and motility before use. Spermatozoa were isolated by the swim-up procedure in HTF or HS solution as previously described (46) and then concentrated by 5 minute centrifugation at ≤500g and supernatant removal.

### Flow Cytometric Assessment of MMP

Human sperm cells were incubated with small molecule uncouplers or vehicles in HS solution for 30 minutes at 37°C, then treated with 50nM MitoTracker Red in the presence of either uncouplers or vehicles for an additional 30 minutes. Subsequently, cells were washed by centrifugation for 5 minutes at 500 g, supernatant was removed, and cells were resuspended in HS solution. Fluorescence was measured by flow cytometry on a BD LSR Fortessa Celeste or BD LSR Fortessa X20, using gates for size and density to exclude debris and aggregates, and a PE-Texas Red Filter set (561nm laser excitation, 610/20 nm emission filter) to record fluorescence (Fig. 1, S1). Fluorescence midpiece location was confirmed by microphotography on an Olympus IX71 inverted microscope using a Hamamatsu Orca-ER digital camera, and a 60x Olympus UPlanSApo objective (Fig. 1). Fluorescence statistics were analyzed using Microsoft Excel version 16 and GraphPad Prism version 9. The subpopulation of cells with the highest gaussian distribution of fluorescence were manually gated and the mean fluorescence of this population was normalized to the average of the means of the same population in the relevant vehicle control samples of each run. Each data point represents the mean fluorescence of the highest normally peak in an individual flow cytometry run of 10,000 cells. Nonlinear least squares regression performed using an unweighted [inhibitor] vs. response three parameter model (Hill Slope = -1.0), identifying “unstable” fits. Outliers tested using a ROUT coefficient of 1%, with no outliers detected. Residuals were checked for normality using the D’Agostino-Pearson omnibus normality test, checked for clustering using the Replicates test, and checked for appropriate weighting using the homoscedasticity test, and passed all tests, except for FCCP which failed the test for homoscedasticity.

### Area Under the Curve Analysis

**F**or all dose-response plots, Area Under the Curve (AUC) analysis was performed using the trapezoid method (ΔX*(Y_1_+Y_2_)/2) on the arithmetic means of the measured parameter at each compound concentration. Each data point represents the area under the curve of all concentrations tested in one biological replicate. Effects of each compound were then compared using Dunnett’s Multiple Comparison’s Test.

### Sperm Motility Assessment

Isolated human sperm were incubated with small molecule uncouplers and vehicles at 37°C for 30 minutes, after which they were mixed with 1% Polyvinyl Alcohol to prevent adherence to glass and loaded into a Leja Standard Count 2-chamber sperm analysis slide (depth 20 micron). Sperm motility was analyzed on a Hamilton-Thorne IVOS I computer-aided sperm analyzer, at 37°C. Motility was measured for least 500 cells in each condition. Progressive motility was defined using Hamilton-Thorne’s proprietary software module. Nonlinear least squares regression performed by GraphPad Prism version 9 using an unweighted [inhibitor] vs. response three parameter model (Hill Slope = -1.0), identifying “unstable” fits. Outliers were tested using a ROUT coefficient of 1% and none were identified. Residuals were checked for normality using the D’Agostino-Pearson omnibus normality test, checked for clustering using the Replicates test, and checked for appropriate weighting using the homoscedasticity test. Residuals for NEN and BAM15 passed all tests, except that BAM15 failed the test for homoscedasticity with a p-value of 0.0207.

Human sperm hyperactivated motility was measured on a Hamilton-Thorne IVOS II computer-aided sperm analyzer, as above, with the definition for hyperactivated motility set as VCL >150 μm/s AND LIN <50% AND ALH >7.0 μm (33).

### Flow Cytometric Assessment of Acrosome Status

Human sperm cells were incubated for 3.5 hours in capacitation media (20% Fetal Bovine Serum and 25mM Sodium Bicarbonate in HS solution) at 37°C and 5% CO_2_. In the final hour of incubation, custom monoclonal antibodies raised in mouse hybridoma cells against the ectodomain of human Izumo1 were added, at a concentration of 10μg/mL. Subsequently 50μM A23187 was added, and cells were incubated at 37°C and 5% CO_2_ for an additional 30 minutes, after which cells were washed by centrifugation 3x (500 g, 5 min) in HS solution, incubated 30 minutes at room temperature with a goat anti-mouse 647 secondary antibody (Invitrogen A32728, final concentration 4μg/mL) in 10% goat serum. After an additional 3 washes by centrifugation, propidium iodide (PI) was added to a final concentration of 1μg/mL, and cells were analyzed by flow cytometry using the same instrumentation as above, except that Izumo-647 was visualized using an APC filter set (bandpass 670/30 or 670/14nm) and Propidium Iodide was visualized using a PE-Texas Red filter set (bandpass 610/20nm). Izumo-647 positive cells were scored as acrosome reacted, and PI negative cells were scored as living.

### Generation of IZUMO1 monoclonal antibody using mouse hybridoma

The IZUMO1 monoclonal antibodies were generated from mouse hybridoma and characterized as previously described (47). Briefly, IZUMO1 antibodies 4E04 and 6F02 belong to the murine IgG1 isotype with kappa light chain. For antibody production, hIZUMO1-positive hybridomas were cultured in medium containing 10% FBS and subsequently adapted to an FBS-free medium. The condition media were harvested, and the IgG antibody were purified by protein G affinity chromatography and by Superdex-200 gel filtration chromatography in a buffer of 150 mM NaCl, 20 mM HEPES pH 7.4. Purified IgG antibodies were concentrated to 1 mg/mL with the addition of 10% (v/v) glycerol for use.

### ATP Assessment

Human Sperm cells were incubated with uncoupler compounds or vehicle for 1 hour at 37°C in white 96-well Costar polystyrene assay plates, under sterile conditions. ATP content was assessed using a PerkinElmer ATPlite Luminescence Assay System Kit, according to manufacturer instructions, and luminescence was imaged on a Biotek Synergy H4 Hybrid plate reader, set to endpoint/kinetic setting (one measurement point per well), with 1 second of integration time and a manual gain setting of 135. ATP content was quantified by interpolating against a standard curve of ATP solutions, per manufacturer instructions. Nonlinear least squares regression performed by GraphPad Prism version 9 using an [inhibitor] vs. response three parameter model (Hill Slope = -1.0), weighted by 1/Y^2^, identifying “ambiguous” fits. Outliers tested using a ROUT coefficient of 1%, with no outliers detected. Residuals were checked for normality using the D’Agostino-Pearson omnibus normality test, checked for clustering using the Replicates test, and checked for appropriate weighting using the homoscedasticity test, and passed all tests.

### Mouse *In Vitro* Fertilization

All mice were kept in the Animal Facility of the University of California, Berkeley, fed standard chow diet (PicoLab Rodent diet 20, #5053, LabDiet), and hyper-chlorinated water *ad libitum* in a room with controlled light (14 hours light, 10 hours darkness) and temperature (23±0.5°C). Animals were humanely killed by CO_2_ asphyxiation and cervical dislocation according to ACUC guidelines with every effort made to minimize suffering.

C57Bl/6N (Charles River) female mice between 4 and 16 weeks old were superovulated by intraperitoneal injection of 5 IU pregnant mare serum gonadotropin (PMSG, Sigma, G4877) and 48 hours later, 5 IU human chorionic gonadotropin (hCG, Millipore, 230734). 13 hours after hCG injection the females were euthanized and cumulus-oocyte complexes were collected from the oviduct ampulla in HTF medium (Embryomax, Specialty Media, Millipore) and incubated for 30 minutes at 37°C and 5% CO_2_ prior to insemination. Simultaneously, mouse sperm were isolated from the cauda epididymis as previously described^36^ and allowed to capacitate at 37°C and 5% CO_2_ for 60-90 minutes in the presence of uncoupler compounds or control vehicle. Sperm and eggs were then mixed to a final concentration of 210,000 sperm/mL in 600μL of HTF medium and incubated at 37°C and 5% CO_2_ for 4 hours. Afterwards, the eggs were washed by mouth pipetting in HTF media to remove excess sperm. A final wash in KSOM media (Zenith Biotech) supplemented with 1mg/mL BSA was then done before dividing the eggs into 10 μL drops of KSOM/BSA media overlaid with embryo-tested light mineral oil (Millipore) for culture at 37°C, 5% CO_2_. Best efforts were made to divide equal numbers of eggs from each mouse into control and uncoupler-treated conditions. 3.5 days following IVF, the number of fertilized eggs was determined visually by assessing the percentage of embryos that reached morula or blastula stage.

### Sperm Whole-Cell Electrophysiology

Whole-cell electrophysiology was performed on isolated human sperm cells as described previously (48,49). Once sperm cell “break in” was achieved in HS solution, cells were subjected to a voltage ramp protocol in the presence of bath and pipette solutions designed to ensure that only H^+^ current across the membrane was measured. Intracellular (pipette) solution contained: 150mM NMDG, 5mM BAPTA, 100mM HEPES, 1mM Tris Chloride, pH 7.5 with MeSO_3._ Extracellular (bath) solution contained: 157mM NMDG, 20mM MES, 1mM MgCl_2_, 100μM ZnCl_2_, pH 6 with MeSO_3_. Compounds and vehicles were applied by perfusion, and currents were recorded until they showed a minimal change between subsequent voltage ramps. Then, compounds were washed out, and subsequent compounds added once currents had returned to baseline values. Representative traces and time courses created in Origin Pro version 9.0.0 and statistical analysis performed in GraphPad Prism 9.

### Mitochondrial Patch-Clamp Electrophysiology

Mouse heart mitochondria were isolated from wildtype and ANT1^-/-^ C57BL/6J2z mice and mitoplasts prepared and patch-clamped as described previously (21). Both the bath and pipette solutions were formulated to record H^+^ currents and contained only salts that dissociate into large anions and cations that are normally impermeant through ion channels or transporters. Pipettes were filled with 130 mM tetramethylammonium hydroxide (TMA), 1.5 mM EGTA, 2 mM Tris chloride, and 100 mM HEPES (or MES). pH was adjusted to 7.5 with d-gluconic acid, and tonicity was adjusted to ∼360 mmol/kg with sucrose. Bath solution contained 100 mM HEPES (or MES) and 1 mM EGTA (pH adjusted to 7.0 with Trizma base, and tonicity adjusted to ∼300 mmol/kg with sucrose).

### Immunocytochemistry

Briefly, sperm were adhered onto poly-D-Lysine coated coverslips, and, in some experiments, incubated 15 minutes at 37°C with 50 nM MitoTracker Red CMXRos, then washed and fixed in 4% PFA for 10 minutes, and washed and additionally fixed in ice-cold methanol for 1 minute. Cells were permeabilized and blocked in 5% BSA and 0.1% Triton X-100 in PBS for 45 minutes, then incubated overnight at 4°C with rabbit polyclonal antibody against Human ANT4 (Novus #NBP1-89074, used at .08 μg/mL) and, in some experiments, a mouse monoclonal antibody against COXIV (COX4l1) (Proteintech #66110-1-Ig, used at 6.8μg/mL). After washing, samples were incubated with secondary antibodies (Jackson Alexa Fluor 647 AffiniPure Donkey Anti-Rabbit IgG (H+L) (code 711-605-152) at a dilution of 1:1000, and Jackson ImmunoResearch anti-mouse Cy3 or Molecular Probes A10521 goat anti-mouse Cy3 at a dilution of 1:1000) for 45 minutes, then washed and mounted using ProLong Gold Antifade mountant with DAPI.

Cells were imaged on an Olympus FV3000 inverted laser scanning confocal microscope, using an Olympus UPLSAPO Super Apochromat 60x oil immersion objective (NA 1.35) and Olympus Z-81226 immersion oil. Laser Scanning was performed by galvanometer, unidirectionally at a rate of 2μs/pixel with no averaging, with pixel sizes of .207 μm/pixel in the x and y direction. Multiple colors were imaged in line-by-line sequential scanning mode, and multiple z-planes were acquired with a thickness of .47 μm, on a linear-encoded stage. Images displayed are maximum intensity z-stacks of all relevant z-planes. All colors were imaged with a pinhole diameter of 233 μM, and laser illumination was passed through a 10% neutral density filter. DAPI was illuminated by a 405 nm laser set to 3% power, and emitted light was passed through a 430-470 nm emission filter and detected with a photomultiplier tube (PMT) set to 500V. Cy3 was illuminated by a 561 nm laser set to 4% power, with a 570-620 nm emission filter and a PMT set to 500V. Alexa 674 was illuminated by a 642 nm laser set to 4% power, with a 650-750 nm emission filter and a PMT set to 550V.

### Super-resolution Microscopy

ICC slides made as described above were imaged using three-dimensional stochastic optical reconstruction microscopy (3D-STORM) (50,51) performed on a home-built set-up using a Nikon CFI Plan Apo λ 100× oil immersion objective (NA 1.45), as described previously (52). In brief, the sample was mounted with an imaging buffer consisting of 5% (w/v) glucose, 100 mM cysteamine, 0.8 mg ml^−1^ glucose oxidase and 40 μg ml^−1^ catalase in a Tris HCl buffer (pH 7.5). For two-color imaging of COXIV and ANT4, the two targets were labeled by Alexa Fluor 647 and CF568, respectively, and were imaged sequentially using 647- and 560-nm excitation lasers. These lasers were passed through an acousto-optic tunable filter and illuminated a few micrometers into the sample at around 2 kW cm^−2^, thus photoswitching most of the labeled dye molecules in the sample into the dark state while allowing a small, random fraction of molecules to emit across the wide-field over different camera frames. Single-molecule emission was passed through a cylindrical lens of focal length 1 m to introduce astigmatism (51), and recorded with an Andor iXon Ultra 897 EM-CCD camera at a frame rate of 110 Hz, for a total of around 50,000 frames per image. Data acquisition used publicly available software (https://github.com/ZhuangLab/storm-control). The raw STORM data were analyzed using Insight3 software (51) according to previously described methods (50,51). Secondary antibodies used: Alexa Fluor 647-labelled goat anti-mouse (Invitrogen, A21236, 1:400) and donkey anti-rabbit (Jackson ImmunoResearch, 711-005-152, 1:70) conjugated with CF568 succinimidyl ester (Biotium, 92131).

### Western Blotting

Human Sperm or HEK293 cells were lysed by addition of RIPA^++^ Buffer (composition: 50mM Tris HCl, pH 7.4; 150mM NaCl; 1% Triton X-100; 0.5% Sodium Deoxylcholate; 0.1% Sodium Dodecyl Sulfate; 1 mM EDTA; 10% glycerol; with Pierce Protease inhibitor tablet or solution added to manufacturer’s instructions), then homogenized by 25 passes through a 21-gauge or smaller syringe, then shook on ice for 30 minutes, and centrifuged at 12000 rpm at 4°C for 20 minutes, after which the supernatant was collected and frozen at -80°C. After thawing, protein concentration was assessed using a Pierce Rapid Gold BCA Protein Assay Kit, per manufacturer instructions, and measured on a Biotek Synergy H4 Hybrid plate reader. A target of 30μg of protein was mixed with 4x Laemmli Buffer and Beta-Mercaptoethanol, heated for 10 minutes at 99°C, then loaded to each well of a precast 4-20% mini-Protean TGX gel (Bio-Rad), and run in an electrophoresis chamber until ladder bands were well-resolved. Transfer onto a PVDF membrane was accomplished with an Invitrogen iBlot Dry Blotting System, following manufacturer instructions, at 20V for 7 minutes. After washing 3x with PBS with 0.1% Tween (PBST), the membrane was blocked in 3% Bovine Serum Albumin for 15-30 minutes, then incubated rocking with primary antibody overnight at 4°C. The following day, the membrane was washed 3x with PBST, and incubated with a 1:15000 dilution of HRP-conjugated Abcam Ab6721 Goat pAb anti-Rabbit IgG secondary antibody for one hour at room temperature. After 3x PBST washes and 2x PBS washes, bands were visualized using a Prometheus ProSignal Pico kit, per instructions, and imaged in a ProteinSimple FluorChem M. Afterwards, images were inverted, rotated, and cropped in ImageJ 1.52a.

## Author Contributions

W.M.S. and P.V.L conceived the project and designed experiments. W.M.S. conducted most experiments and analyzed most data, except as noted below, and wrote the manuscript. N.T.P. and W.M.S. performed *in vitro* fertilization experiments. B.U. and W.M.S. conducted STORM and B.U. analyzed data, under supervision of K.X. S.T. produced Izumo1 antibodies used during acrosome reaction assays. E.T. and J.L. assisted with multiple experiments. L.J. and J.S. enabled access to and training on computer-aided sperm analysis machines for motility assays performed by W.M.S. A.M.B. performed mitochondrial patch-clamp experiments and analyzed that data, under supervision of Y.K. P.V.L and W.M.S. performed human sperm patch-clamp electrophysiology. All of the authors discussed the results and commented on the manuscript.

Conceptualization: W.M.S and P.V.L

Data Curation: W.M.S.

Formal Analysis: W.M.S. and A.M.B

Funding Acquisition: P.V.L. and W.M.S.

Investigation: W.M.S., N.T.P., B.U., E.T., J.L., A.M.B., P.V.L

Methodology: W.M.S., B.U., A.M.B, P.V.L

Project Administration: W.M.S.

Resources: W.M.S., S.T., L.J., J.S., K.X., Y.K., P.V.L.

Supervision: J.S., K.X., Y.K., P.V.L. Validation: W.M.S.

Visualization: W.M.S., B.U., A.M.B

Writing – Original Draft Preparation: W.M.S.

Writing – Review & Editing: All Authors

## Acknowledgments

This work was furthered by the support and guidance of Dr. Denise Schichnes, using the equipment of the CNR Biological Imaging Facility at the University of California, Berkeley. Research reported in this publication was supported in part by the National Institutes of Health S10 program under award number 1S10OD018136-01. The content is solely the responsibility of the authors and does not necessarily represent the official views of the National Institute of Health. The authors also wish to thank Dr. Liliya Gabelev Khasin, Dr. Caroline Williams, and Dr. Lisa Treidel, and Dr. Erwin Goldberg for their assistance with this project.

This work was supported by an Irving H. Wiesenfeld Fellowship administered by the UC Berkeley Center for Emerging and Neglected Diseases to W.M.S., a Male Contraceptive Initiative Graduate Fellowship to P.V.L. and Liliya Gabelev Khasin, NIH NICHD grant K99HD104924 to S.T., NIGMS grant R35GM136415 to Y.K., Pew Biomedical Scholars Award to P.V.L. and K.X. and Bakar Fellow Spark Award to P.V.L. This material is based upon work supported by the National Science Foundation Graduate Research Fellowship Program, given to W.M.S. under grant numbers DGE 1752814 and DGE 2146752. Any opinions, findings, and conclusions or recommendations expressed in this material are those of the author(s) and do not necessarily reflect the views of the National Science Foundation.

## Abbreviations

ANT: Adenine nucleotide translocase
ALH: Amplitude of lateral head displacement
ATP: Adenosine triphosphate
AUC: Area under the curve
BAM15: (N5,N6-bis(2-Fluorophenyl)-[1,2,5]oxadiazolo[3,4-b]pyrazine-5,6-diamine)
BAPTA: (1,2-bis(o-aminophenoxy)ethane-N,N,N′,N′-tetraacetic acid)
BSA: Bovine serum albumin
CATR: Carboxyatractyloside
COXIV: Cytochrome c oxidase subunit 4 isoform 1, mitochondrial
DNP: 2, 4, Dinitrophenol
EGTA: (ethylene glycol-bis(β-aminoethyl ether)-N,N,N′,N′-tetraacetic acid)
FCCP: Trifluoromethoxyphenylhydrazone
FDA: Food and Drug Administration
HEPES: (4-(2-hydroxyethyl)-1-piperazineethanesulfonic acid)
HRP: Horseradish peroxidase
HS: High Saline
HTF: Human Tubal Fluid
ICC: Immunocytochemistry
IMM: Inner mitochondrial membrane
KSOM: Potassium-supplemented simplex optimized medium
LIN: Linearity
MES: 2-(N-morpholino)ethanesulfonate.
MMP: Mitochondrial membrane potential
NEN: Niclosamide Ethanolamine
NMDG: N-methyl-D-glucamine
PI: propidium iodide
PMT: Photomultiplier tube
PVDF: polyvinylidene difluoride
STORM: Stochastic optical reconstruction microscopy
Tris: tris(hydroxymethyl)aminomethane
VCL: Curvilinear velocity

---

**Figure S1.**
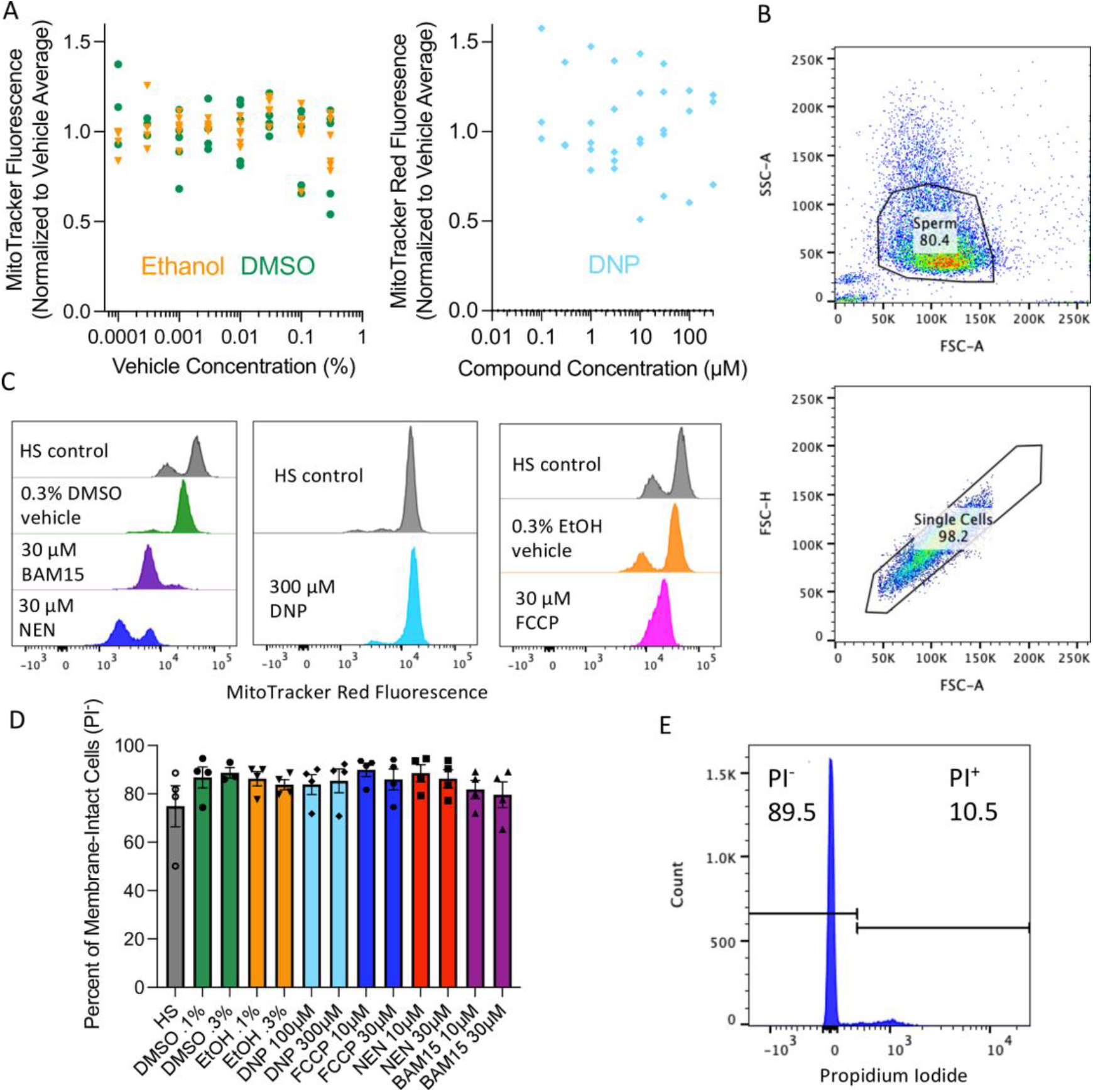
Validation of flow cytometric assessment of human sperm mitochondrial membrane potential. **A) Left:** Effect of compound vehicles on MitoTracker fluorescence. **Right:** Effect of the uncoupler 2, 4, DNP on MitoTracker Fluorescence. In both, nonlinear regression performed, but not shown or reported because adjusted R^2^ values were <.1. B) Representative gating logic for flow cytometry experiments, isolating individual sperm cells from debris (top) and aggregations (bottom). C) Representative flow cytometry histograms for MitoTracker Red CMX-Ros Mitochondrial Membrane Potential Assays. D) Percent of propidium iodide-excluding cells (membrane intact) after 30-minute incubation with compounds, as measured by flow cytometry. Each data point represents the average of 10,000 cells, and error bars represent SEM. All conditions were compared to DMSO .1% using Dunnett’s Multiple Comparisons Test, and no conditions were significantly different. E) Representative flow cytometry result of propidium iodide vitality assay.

**Figure S2:**
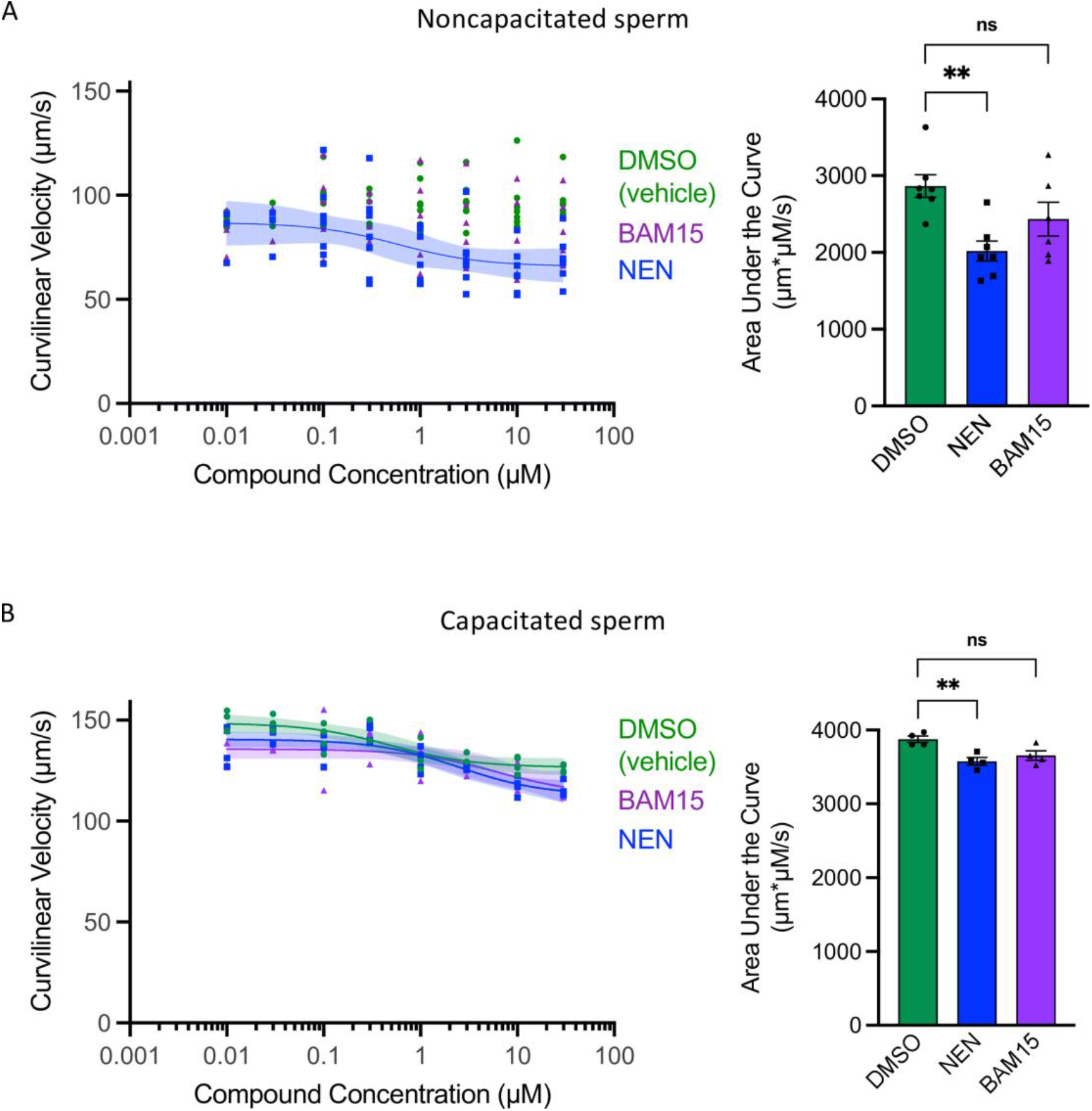
Effects of small molecule uncouplers on human sperm curvilinear velocity as measured by computer-aided sperm analysis. **A)** Curvilinear velocity analysis of noncapacitated human sperm cells. Left: Each data point represents the average curvilinear velocity of at least 200 sperm cells. Nonlinear least squares regression performed using an unweighted [inhibitor] vs. response three parameter model (Hill Slope = -1.0), identifying “unstable” fits. Shaded areas indicate 95% confidence bands for true location of each nonlinear fit curve. The nonlinear regression fit for NEN had an Adjusted R^2^ value of 0.1833. BAM15 & DMSO nonlinear regressions not shown because their adjusted R^2^ values were below 0.1. NEN IC_50_ Asymmetrical 95% CI interval: 0.07433 μM to 6.950 μM. Outliers tested using a ROUT coefficient of 1%, with two outliers in the DMSO condition excluded. NEN residuals passed the Replicates and homoscedasticity tests, but failed the D’Agostino-Pearson omnibus normality test with a p-value of .0366. Right: Area Under the Curve analysis for data at left. Each data point represents the area under the curve of one biological replicate. **B)** Curvilinear Velocity analysis of capacitated human sperm cells. Left: Nonlinear Regression performed as in A. The nonlinear regression fit for DMSO had an Adjusted R^2^ value of 0.7328 with an IC_50_ 95% Confidence Interval of 0.1126 μM to 1.925 μM. The nonlinear regression fit for NEN had an Adjusted R^2^ value of 0.7275 with an IC_50_ 95% Confidence Interval of 0.9469 μM to 7.230 μM. The nonlinear regression fit for BAM15 had an Adjusted R^2^ value of 0.3885 with an IC_50_ 95% Confidence Interval of 0.8117 μM to 294309381 μM. All tests performed as in A, with no outliers detected, and with residuals passing all tests, except that the residuals for NEN failed the test for homoscedasticity with a P-value of .006. Right: Area Under the Curve analysis for data at left.

**Figure S3:**
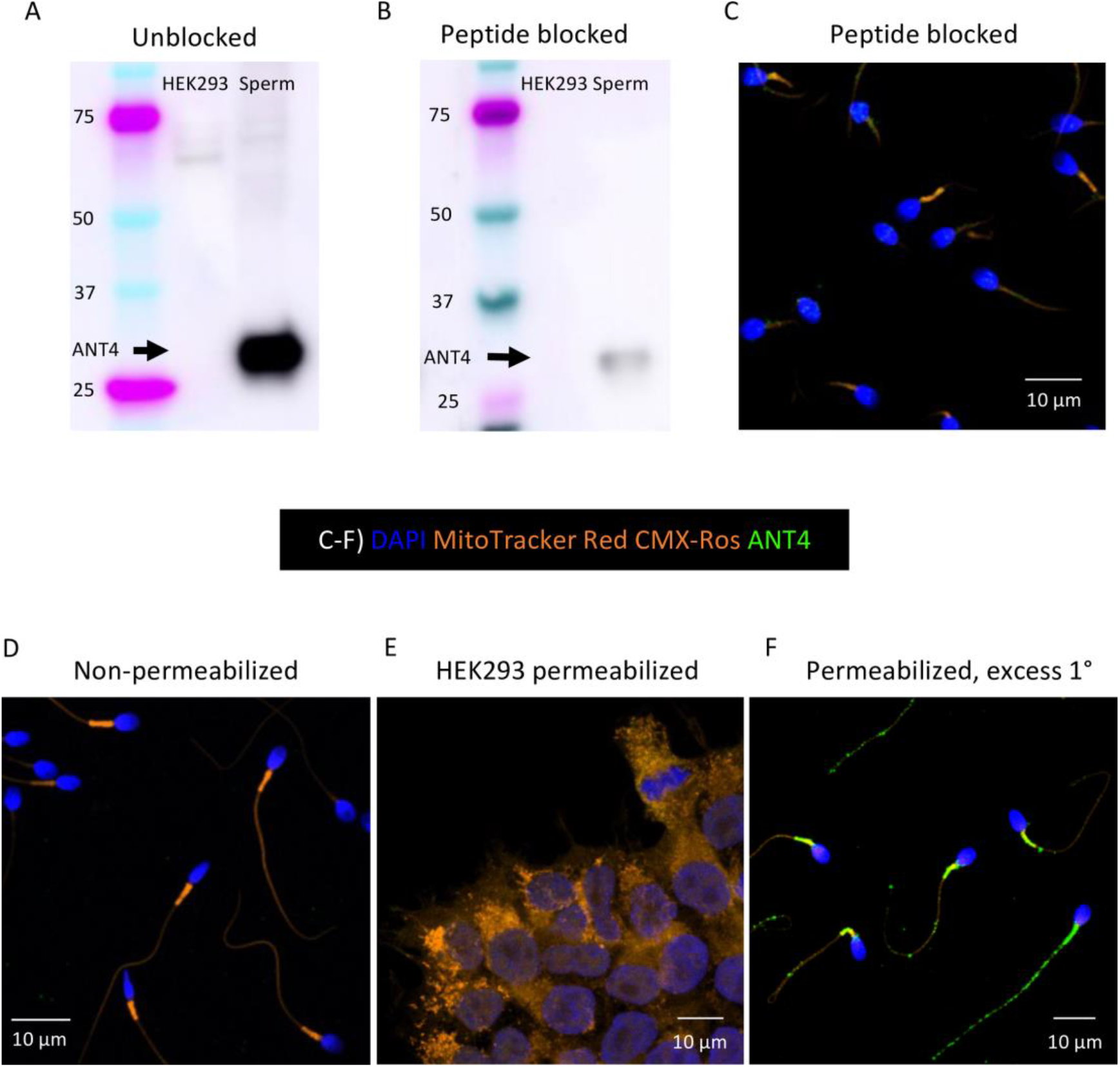
Human ANT4 polyclonal antibody validation. **A**,**B)** hANT4 Western blot of HEK293 cells and human sperm. A), unblocked; B) blocked with immunizing peptide. C) ICC of permeabilized human sperm incubated with hANT4 pAb preblocked with immunizing peptide and 50 nM Mitotracker Red CMX-Ros. **D)** ICC of non-permeabilized human sperm labelled as in C. E) ICC of permeabilized HEK293 cells labelled as in C. F) Permeabilized human sperm cells incubated with excess hANT4 pAb primary antibody (.4 μg/mL) and 50 nM MitoTracker Red CMX-Ros. In C-F, Orange – MitoTracker Red CMX-Ros, Blue – Dapi, Green – hANT4. All images are maximum intensity projections of sequential confocal image z-slices.

**Figure S4:**
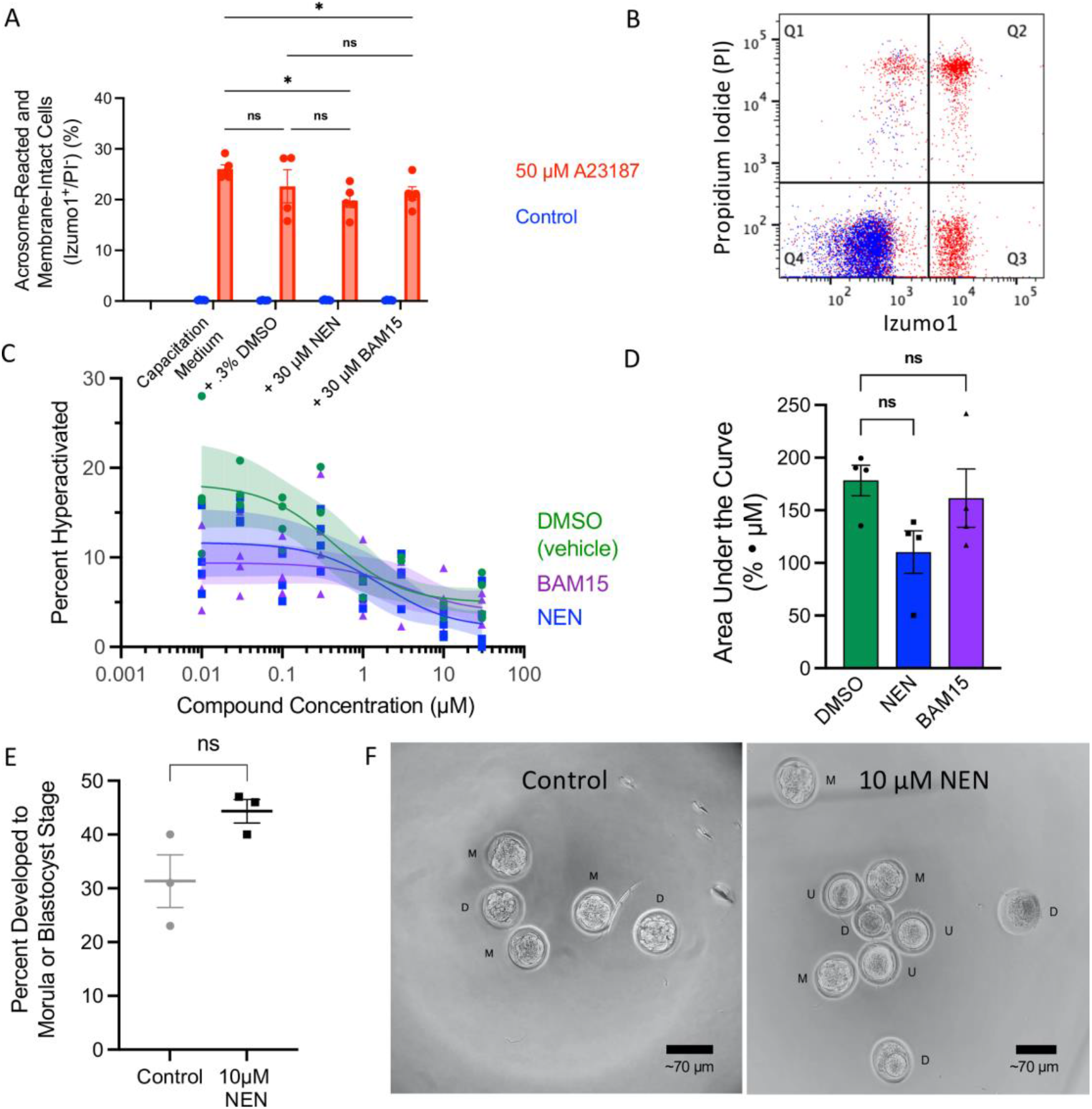
NEN and BAM15 have no effect on human sperm acrosome reaction competence or human sperm hyperactivated motility, and NEN has no effect on Mouse IVF success. **A)** Percentage of unfixed, capacitated human sperm cells that were acrosome-reacted and membrane-intact (Izumo1^+^/Propidium Iodide^-^) as measured by flow cytometry, after incubation with and without the calcium ionophore A23187. Each data point represents the percentage of 100,000 cells showing this fluorescence pattern. Error bars represent SEM, and significance was calculated by Tukey’s multiple comparisons test, with individual variances computed for each comparison. All combinations were compared, but for clarity, not all comparisons are shown. All non-shown comparisons were not significant. **B)** Representative data plots from one run of flow cytometry, comparing ionophore A23187-treated (red) and control cells (blue). **C)** Representative confocal microscopy image of A23187-treated sperm cells taken from flow cytometry run, showing Izumo1 staining in red, and DAPI staining in blue. AR: Acrosome Reacted, AI: Acrosome Intact. **D)** Percent of capacitated human sperm cells displaying hyperactivated motility, as measured by computer-aided sperm analysis. Each data point represents the percent of at least 200 sperm cells showing hyperactivated motility. Nonlinear least squares regression performed using an unweighted [inhibitor] vs. response three parameter model (Hill Slope = -1.0), identifying “unstable” fits. Shaded areas indicate 95% confidence bands for true location of each nonlinear fit curve. The nonlinear regression fit for DMSO had an Adjusted R^2^ value of 0.6501, and an IC_50_ 95% CI of 0.1052 μM to 2.131 μM. The nonlinear regression fit for NEN had an Adjusted R^2^ value of 0.5062, and an IC_50_ 95% CI of 0.2957 μM to 12.04 μM. The nonlinear regression fit for BAM15 had an Adjusted R^2^ value of 0.1667, and an IC_50_ 95% CI of 0.1418 μM to +infinity. **E)** Area under the curve analysis of hyperactivation data from panel E. Each data point represents the area under the curve of one biological replicate, error bars represent SEM, and statistical significance calculated using Dunnett’s Multiple Comparisons Test. **F)** Quantification of mouse eggs that developed to morula or blastocyst stage after *In vitro* fertilization (IVF) using SMU-treated sperm. Significance calculated using a one-tailed ratio paired t-test. **G)** Representative micrographs of mouse embryos 3.5 days after IVF. Left, oocytes exposed to control sperm; Right, oocytes exposed to 10 μM NEN-treated sperm. M - Morula, D - Dead, U - Unfertilized.

**Figure S5:**
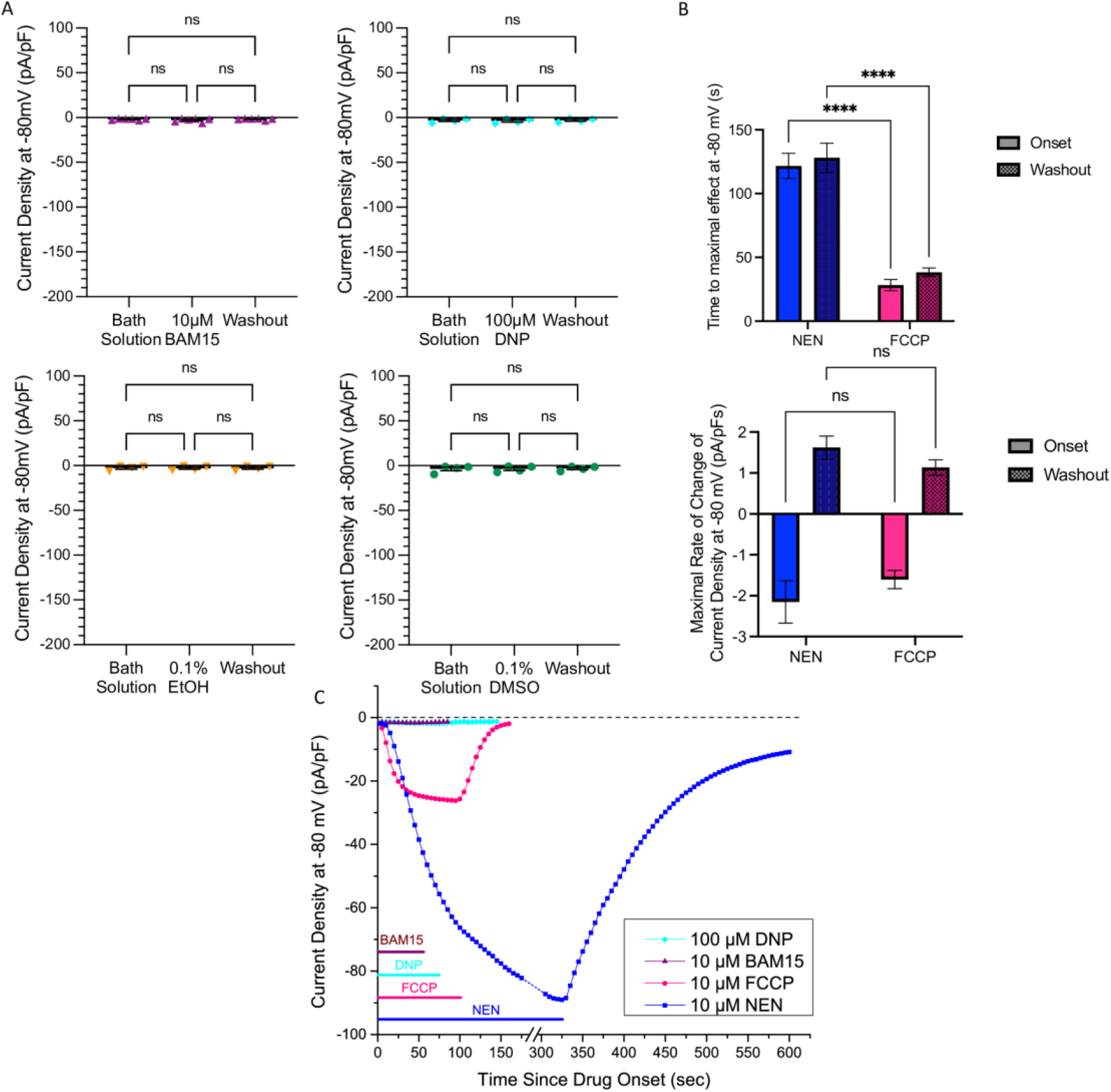
Supplemental electrophysiology quantifications. **A)** 10 μM BAM15, 100 μM DNP, 0.1% EtOH, and 0.1% DMSO have no effect on sperm plasma membrane current densities. Significance calculated using Tukey’s multiple comparisons test. **B)** NEN takes longer to reach maximal uncoupling than FCCP (top), but their maximum rates of current density increase are similar (bottom). Significance calculated using Šídák’s multiple comparison’s test, with a single pooled variance **C)** Time courses of current density at -80 mV during application of compounds. Colored bars represent time period during which each compound was being applied. Dotted line in NEN time course represents a period where other electrophysiological tests were performed on that cell.

